# AI platform for CRISPR functional mapping and function-based drug design

**DOI:** 10.64898/2026.05.06.722817

**Authors:** Jason C. Ngo, Vivien A.C. Schoonenberg, Renu Nandakumar, Xuebing Wu, Falak Sher

## Abstract

Conventional structure-based drug design has high clinical failure rates due to the disconnect where binding affinity does not guarantee safe functional modulation. To bridge this gap, we present CRISPRtile, a cloud-based platform for function-based drug design. By deriving library coverage optimization equations and leveraging AI to correct CRISPR guide biases, we generated toxicity and functional landscapes with over threefold error reduction compared to conventional methods. These maps bypass error and orders of magnitude higher computational cost in structure-based pipelines by enabling AI prediction of drug interaction directly from nontoxic functional sequences, while predicting brain penetration with benchmark leading performance. We demonstrate CRISPRtile by mapping the NLRP3 inflammasome and identifying FDA approved drugs with previously unrecognized ability to modulate it, revealing strategies to amplify or inhibit our immune response to homeostatic perturbations. These advances establish a generalizable strategy for the systematic discovery of safe functional modulators.

## Introduction

Virtual screening offers a cost-effective way to search the therapeutic space, yet lacks direct assessment of toxicity or function^1^. Variant pathogenicity predictions are insufficient for identifying druggable regions, as they do not disentangle toxicity from loss of function. Population frequency^2^ can not be used because mutations can be recent, with a low contribution to the maximum filial generation number that selection pressure has not been placed on it to reflect function. Consequently, drug discovery has a disconnect between binding and functional modulation, explaining why ∼90% fail clinical trials, ∼40-50% from functional efficiency, ∼30% from toxicity, and ∼10-15% from delivery^3^.

Developing nontoxic modulators requires understanding which residues drive function or toxicity, a challenge that CRISPR struggles to address with sufficient resolution^4-14^. While precise editors^5-7^ enable functional mapping, they are constrained by substitution capabilities^6,7^ and guide dependent biases^8,9^, due to differences in editing efficiency and indel profiles. Benjamini–Hochberg does not consider the correlated effects of substitutions, leading to false positives^10^. Conservative substitutions can mask essential residues, while gain-of-function mutations can exaggerate nonessential residues.

To overcome these limitations, we present CRISPRtile, which leverages Cas9^11,12^ saturation mutagenesis^13,14^ to expand the mutational space at high efficacy and avoid the problems that come with substitutions^5,10^. CRISPRtile resolves the signal to noise problems in CRISPR screens^8,9^ by optimizing library coverage and correcting for guide dependent biases with AI. This reduces error and enables the construction of high fidelity maps of toxicity and function that are the foundation for predicting drug interactions.

We demonstrate this on NLRP3, a driver of IL-1β and IL-18 in response to homeostatic perturbations to combat malignancies and pathogens^16-19^. This response induces DNA double strand breaks in bystander cells^20^, accelerating epigenetic erosion and compromising tissue homeostasis^21^. This dichotomy is evident in the *APOE4* polymorphism, which confers survival advantages in melanoma^22^, but shows heightened NLRP3 activity^23^ that exacerbates neurodegeneration^24^ in Alzheimer’s Disease through inflammatory responses to sterile stressors, such as environmental microplastics^25,26^. This suggests that the tradeoff of being able to sense unspecific homeostatic perturbations to target malignancies and pathogens that may have evolved to avoid specific targeting is the cost of being unable to distinguish from sterile stressors, limiting NLRP3 activation and highlighting a need for context dependent modulation.

The problem is that NLRP3 inhibitors failed clinical trials due to toxicity^27^ and there are no NLRP3 amplifiers that avoid basal activation. Using CRISPRtile, we identify FDA approved drugs with established safety profiles and previously unrecognized ability to inhibit or amplify NLRP3 without inducing basal activation, overcoming safety bottlenecks and demonstrating CRISPRtile’s potential for function-based drug design.

## Results

### CRISPRtile optimizes library sampling and gating for pooled CRISPR screening

Pooled CRISPR screens target a multiplicity of infection (MOI) between 0.3-0.5, with coverage of 25–1,000 cells per guide^28^. Uniform library representation is assumed, but guide distributions often span five orders of magnitude^29^. Low cell numbers introduce stochastic noise, while excessive cell numbers are costly and add noise from sampling bias. Virion amount also matters because if the abundance of a guide is low, small differences in abundance affect the final fold change. Conversely, if the abundance of a guide is high, the cells are overtransduced, confounding signals with multi-guide effects. To resolve this, we derived equations to optimize the cell and virions required to satisfy coverage thresholds given guide frequencies for *k* guides per cell, allowing for combinatorial screens requiring multi-guide combinations. We model the acquisition of the library through the weighted coupon collector problem where an upper bound for the number of draws to obtain all coupons each with a probability *p* where 0 < *p* < 1 is given by^30^

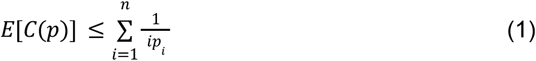

where *p*_1_ ≤ *p*_2_ … *p*_*n*_. Adding our guide frequencies

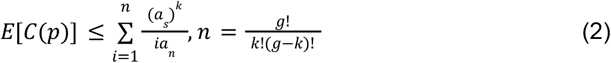

where *a*_*s*_ is the sum abundance of all guides, *a*_*n*_ is the product of the nth guide combination abundances, *g* is the number of different guides we have in our library and *k* is the number of guides we want in a cell. The fraction of cells we have that have k guides *f* is given by the poisson distribution

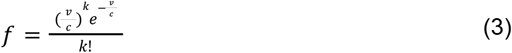

where *v* is the number of effective virions and *c* is the number of cells in our screen. Combining equation (2) and equation (3) to get the number of cells in our screen *c*

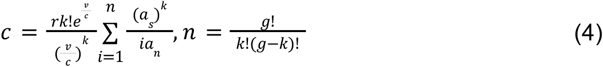

where *r* is the number of single cell replicates we want in our screen. After taking the derivative of equation (3) with respect to *v*

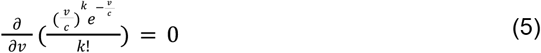

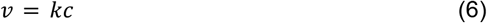

the amount of effective virions *v* to add that maximizes the fraction of cells with k virions is just *kc*. Substituting equation (6) into equation (4) and get

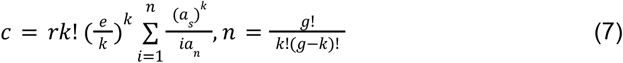

To calculate the number of effective virions *v* we have added to the cells given the number of cells that are positive for infection in the titer *p*, and the initial number of cells in the titer *c*_*i*_, we have the binomial distribution probability mass function where:

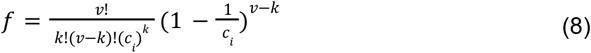

We set *k* = 0 to calculate the fraction of cells with no guides and we get:

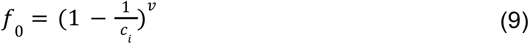

We solve for *v* and we get the number of effective virions in our titer:

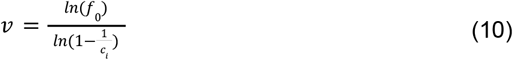

So the volume of titer to add to a screen *A* and upper limit if we want 1 guide per cell where *L* is the volume we used for the titer would be:

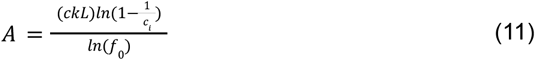

By replacing heuristic-based transduction that does not consider guide frequencies with a mathematical framework, CRISPRtile leverages the guide frequencies to establish a starting baseline, maximizing library representation and mitigating sampling errors in skewed distributions.

To overcome the subjectivity and dimensionality limitations of manual gating in Fluorescence Activated Cell Sorting (FACS), we developed the CRISPRtile FACS module, which replaces heuristic boundary setting with an automated data driven strategy for classification. We used the module to optimize the NLRP3 activation protocol and sorting conditions. Utilizing ASC specks as a direct readout for NLRP3 activation^16^, CRISPRtile distinguishes activated from unactivated populations (Fig. 1a). The robustness of this gating strategy was confirmed by resorting, which demonstrated high purity (Fig. 1a, Supplementary Fig. S1). To interrogate NLRP3, we engineered a tiling library targeting *NLRP3* and *ASC*, coupled with specificity controls, including sgRNAs tiling alternative proteins involved in inflammasome activation (*NLRP1*^16^, *IFI16*^*31*^, *NLRC4*^16^, *AIM2*^16^, *MEFV*^32^, *PLCG2*^33^), inflammatory caspases (*CASP1*^16^, *CASP4*^34^, *CASP5*^34^), as well as negative control guides^35^ to ensure NLRP3 dependent readout (Fig. 1b). We observed differences in dropout scores calculated by the log2 fold change in read counts at day 14 over day 6 across the targets (Kruskal-Wallis H=256.0, p<0.001), with *NLRP3* targeting guides exhibiting an 11.5% reduction in the median day 14 read count over day 6 read count relative to controls (p<0.001, Fig. 1c). The screen detected differences in functional scores calculated by the log2 fold chance in read counts of the activated over unactivated group (Kruskal-Wallis H=1226.4, p<0.001, Fig. 1d). Guides targeting *NLRP3* and *ASC* lowered median activated over unactivated fraction by 85.9% and 97.0%, respectively, compared to controls (p<0.001 for both, Fig. 1d). In contrast, no significance was detected in all other targets, confirming the assay’s specificity for NLRP3 activation (Fig. 1d).

**Figure 1.**
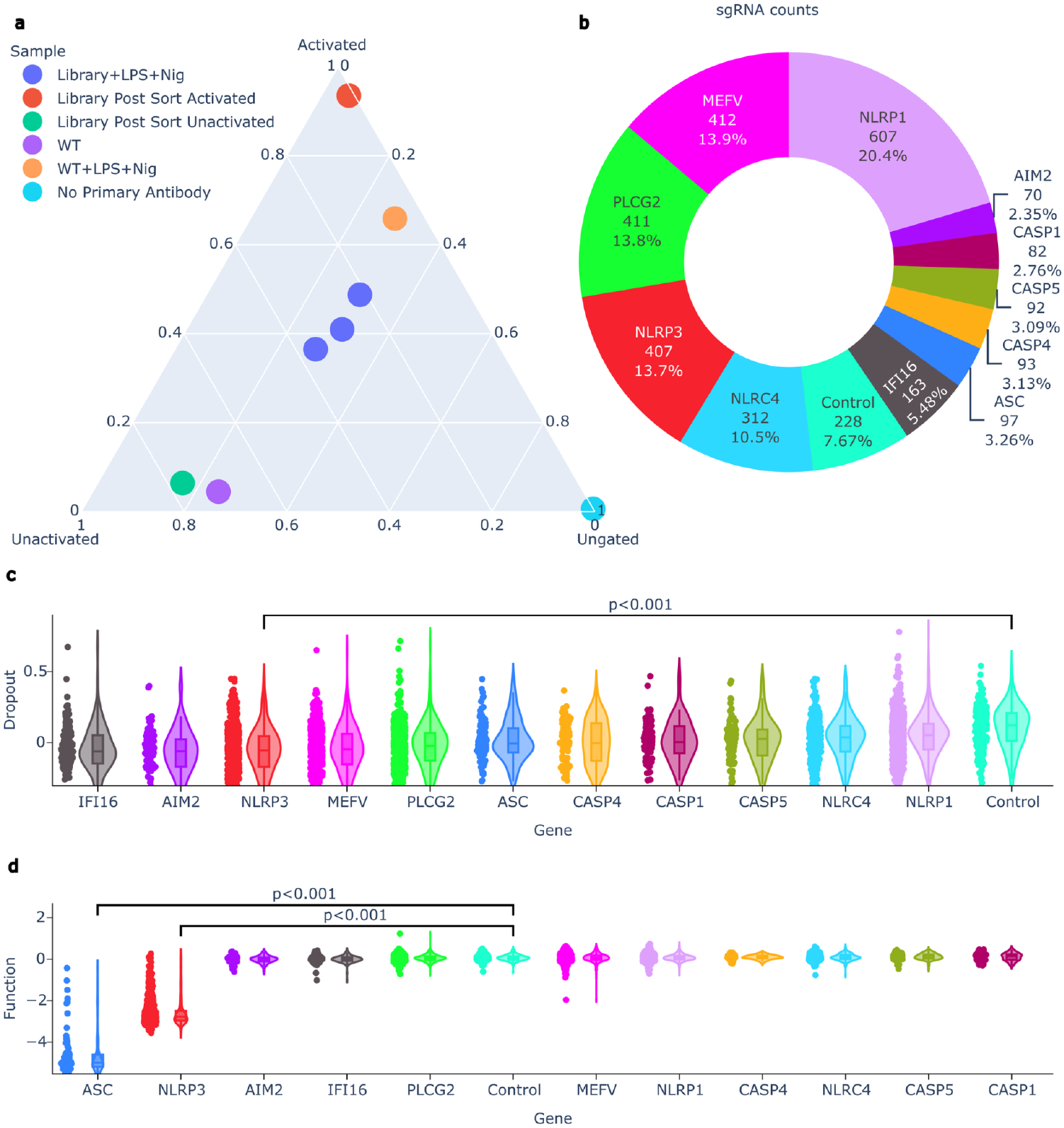
setup of CRISPRtile and the pooled CRISPR screen. **a**, CRISPRtile FACS module separation of activated, unactivated, and unspecific cells for the pooled CRISPR screen by fraction of cells in each group. **b**, Representation of sgRNA library distribution for the pooled screen. **c**, Dropout scores, defined by log2 fold change in read counts at day 14 over day 6. Targets are stratified by gene and ranked by median dropout score **d**, Function scores, defined as the log2 fold change of read counts in the activated population over unactivated population. Targets are stratified by gene and ranked by median function score. For **c** and **d**, P values were determined using a Kruskal-Wallis test (H=256 and 1226.4 respectively) followed by Dunn’s post-hoc test with Bonferroni correction for multiple comparisons. Box plots indicate median (bounds), 25th and 75^th^ percentiles (bounds), and 1.5xIQR (whiskers). Individual data points represents the mean log2 fold change across n = 3 biological replicates for unique sgRNAs (NLRP3, n=387, ASC, n=93, Controls, n=215)

### CRISPRtile achieves superior predictive accuracy and biological concordance

Standard analytical methods for CRISPR tiling screens rely on experimental readouts, which are confounded by guide dependent bias and are unable to comprehensively map the dropout and functional landscape due to limited targeting capabilities. To deconvolute signal from these confounders, CRISPRtile uses machine learning, trained to predict experimental readouts from features such as guide dependent scores^8,9^, 3D amino acid positioning^1^, PROVEAN conservation scores^3^, protein disorder^36^, Solvent-Accessible Surface Area^1,37^, and amino acid identity^1^.

To evaluate the model’s capacity to capture the dropout and functional landscape prior to bias correction, we used 8-fold cross-validation. By evaluating performance of out-of-fold (OOF) predictions, where each experimental data point is scored by a sub model blind to that point during its fitting phase, we ensure a stringent internal validation without sacrificing training volume. CRISPRtile demonstrated predictive fidelity against conventional methods on the uncorrected OOF data (Dropout RMSE=0.05, Function RMSE=0.21), outperforming ProTiler^38^ (Dropout RMSE=0.29, Function RMSE=0.77) and CRISPRO^36^ (Dropout RMSE=0.28, Function RMSE=0.72, Fig. 2b). This confirms the model learned generalizable rules for how guide dependent scores affect the dropout and functional landscape to yield the experimental readout. Furthermore, OOF predictions exhibited near linear proportionality to the experimental ground truth, achieving exceptional Pearson correlations (r=0.990 for Dropout, r=0.986 for Function, p<0.001) and high rank order preservation (Rho=0.967 and 0.951 respectively, p<0.001), outperforming ProTiler^38^ and CRISPRO^36^ (Fig. 2b).

**Figure 2.**
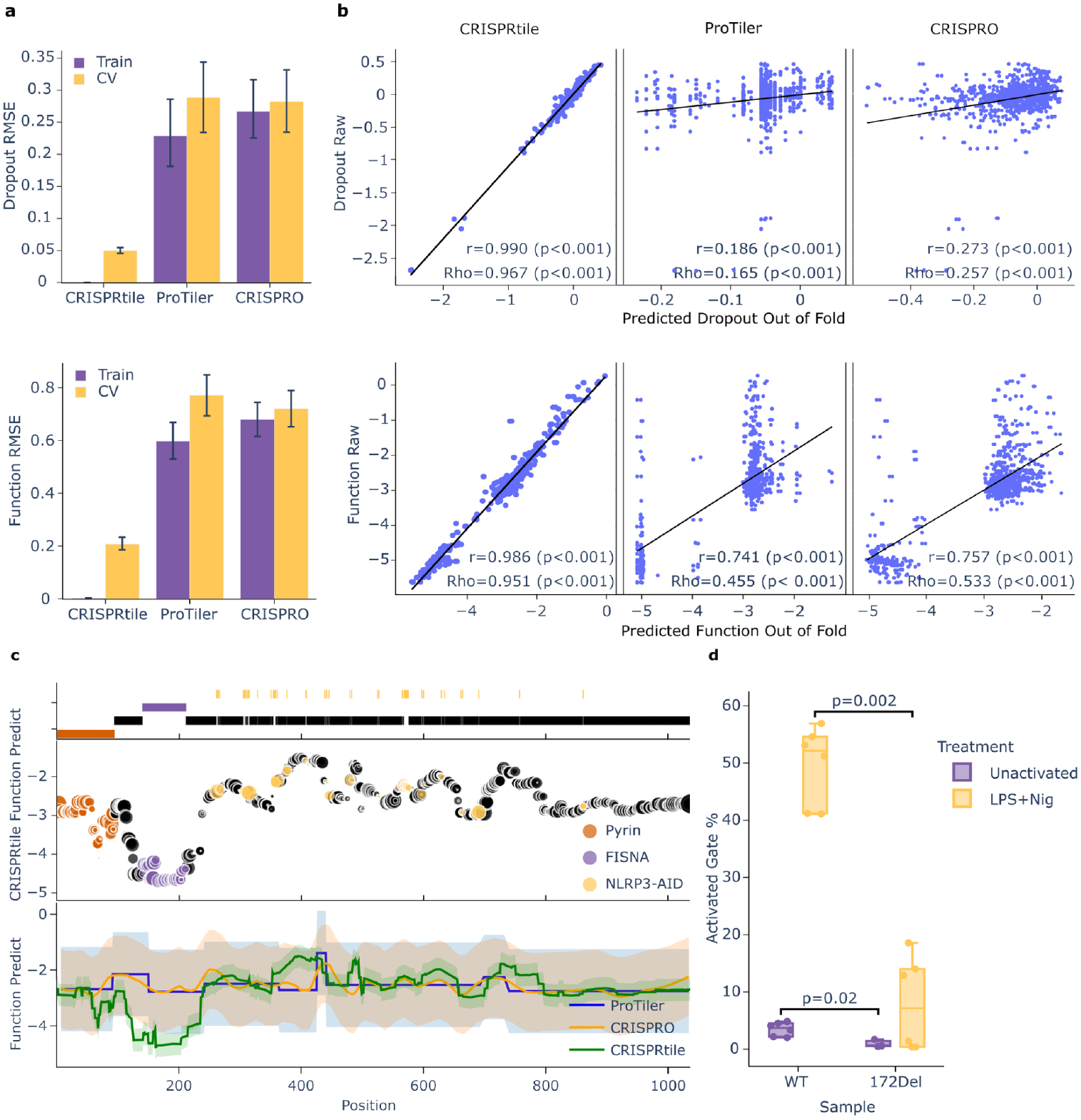
Benchmarking and validation of CRISPRtile. **a**, RMSE of the dropout and function score of CRISPRtile compared to conventional methods. Data is the median and 95% confidence interval of the median from bootstrapping 10,000 times. The train label is performance when trained across entire dataset and cross-validation label is performance on out-of-fold amino acid positions not used for training through 8-fold cross-validation (n = 960 amino acid positions from the mean of n = 3 biological replicates) **b**, Predicted scores on out-of-fold data from 8-fold cross-validation plotted against actual scores. (n=960 amino acid positions from the mean of n = 3 biological replicates) P values were determined using a two-sided Pearson and two-sided Spearman test. **c**, Mapping of pathogenic variants and functional domains of NLRP3 on CRISPRtile predictions along with conventional methods. The size of the points represents the Solvent-Accessible Surface Area. The shaded region represents the 95% prediction interval. **d**, FACS of WT (n = 6 biological replicates), WT+LPS+Nigericin (n = 6 biological replicates), NLRP3 172del clone (n = 3 biological replicates), and NLRP3 172del clone+LPS+Nigericin (n = 6 biological replicates) on ASC speck formation gated by FACS module of CRISPRtile. P values were determined using a two-sided Mann-Whitney U exact test.

While our internal OOF metrics confirms the model captures the raw experimental landscape, our primary objective is to isolate unbiased biological significance. CRISPRtile normalizes the guide dependent features to a global median and predicts a corrected score that decouples guide bias from true functional and toxicity signals, enabling high resolution comparisons across sequences. To validate the accuracy and generalizability of these predictions, we evaluated the model against an independent test set, excluded from all prior phases of model training, from external data composed of clinical variants and annotated regions (Fig. 2c-d, Supplementary Fig. S2f).

Given that regions in NLRP3 are well characterized, NLRP3 serves as a rigorous benchmark for CRISPRtile’s ability to resolve critical regions and clinically relevant hotspots with high precision. CRISPRtile identified known functional domains characterized by UniProt^39^, with signal troughs corresponding to functional regions in the Pyrin domain, responsible for binding to ASC^40^, and the FISNA domain, a sensor required for ASC speck formation^41^, both required for activation in WT (Fig. 2c). We overlaid all pathogenic and likely pathogenic NLRP3 variants from Infevers^15^ (n=46 residues) known to cause autoinflammatory disease from basal activation of NLRP3 onto our CRISPRtile predictions and observed concordance, where mutations that cause basal activation localized to our functional peaks (Fig. 2c). Furthermore, we observed a flat region after NLRP3 AA800, consistent with studies showing low density of pathogenic variants in this tail^15^, low ASC speck formation from mutations in this region^42^, and NLRP3 AA1-686 as a minimum truncated variant required for activation^43^ (Fig. 2c).

To experimentally validate a region uniquely resolved by CRISPRtile, we generated a single amino acid deletion at NLRP3 AA172, a signal not found in raw data nor in ProTiler^38^ or CRISPRO^36^ predictions (Fig. 2c, Supplementary Fig. S2a-e). In agreement with CRISPRtile, this ablation reduced mean activated gate % on ASC specks, a measure of NLRP3 activation, compared to WT in both unactivated and LPS+Nig treatment by 74.0% (p=0.02) and 81.4% (p=0.002) respectively (Fig. 2d), consistent with studies showing reduced ASC speck formation when part of the FISNA region is deleted^41^, and confirming the platform’s ability to uncover functional sequences missed by ProTiler^38^ and CRISPRO^36^.

To evaluate CRISPRtile’s ability to function on diverse datasets, we applied it to train a new model on an external dataset for fetal hemoglobin (HbF) induction^14^. Querying ClinVar^44^ for HbF inducing variants within these tiled genes yielded three missense mutations in ZBTB7A. CRISPRtile outperformed the original CRISPRO^36^ analysis by correctly ranking the phenotypic magnitudes of these variants (Supplementary Fig. S2f). C384W, which drives 100% HbF induction^45^, localized to the apex of the CRISPRtile predictive peak (Supplementary Fig. S3f). In contrast, CRISPRO^36^ erroneously predicted C384W to have the lowest HbF induction among the three variants (Supplementary Fig. S3f). Furthermore, CRISPRtile avoided a false positive prediction for D452N^46^. While CRISPRO^36^ mapped D452N to its highest peak, CRISPRtile mapped it within a non-peak, consistent with the variant’s normal level of 0.5% HbF^46^, despite being a pathogenic, nonconservative mutation (Supplementary Fig. S3f). These findings demonstrate the capacity of CRISPRtile to extract robust, clinically actionable insights from diverse tiling screens.

### CRISPRtile deconvolutes guide dependent bias from biological signals to reveal functional sequences

To evaluate CRISPRtile against established metrics, we compared our functional maps with Provean^2^ and Disorder^36^ scores (Fig. 3a, Supplementary Fig. S3). While PROVEAN classifies activating and deactivating mutations as deleterious, it lacks the directional resolution to distinguish between opposing phenotypes in NLRP3 (Fig. 3a). Furthermore, the functional region identified by CRISPRtile exhibits high disorder, presenting a historical barrier to structure-based drug design (Fig. 3a). To determine if CRISPRtile could identify targetable vulnerabilities, we mapped our functional scores onto the AlphaFold structure of NLRP3 and observed that functionally indispensable residues colocalize and are solvent exposed (Fig. 3b). Projecting these scores onto the cryo-EM structure of compound bound, unactivated NLRP3 (7PZC, Fig. 3c)^47^, CRISPRtile delineated the binding sites for ADP at NLRP3 AA231-234 (Fig. 3d) and MCC950 at NLRP3 AA228 (Fig. 3e), solely from CRISPRtile functional maps.

**Figure 3.**
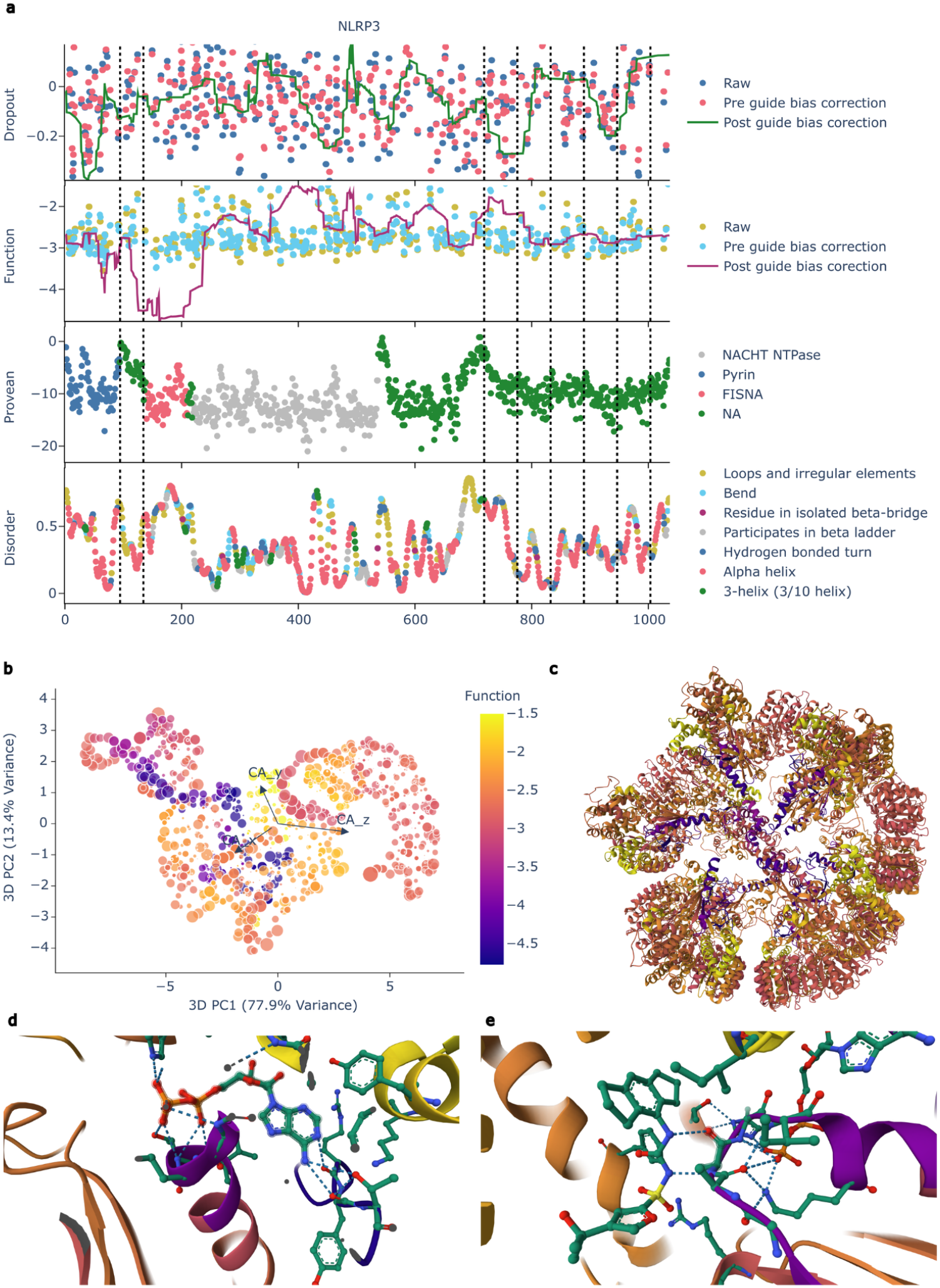
Mapping of CRISPRtile. **a**, CRISPRtile map of NLRP3 by amino acid position where exons are separated by black dotted lines. Raw data points represents the mean log2 fold change across n = 3 biological replicates for unique sgRNAs (NLRP3, n = 387). The pre guide bias correction is the out-of-fold prediction from 8-fold cross-validation and post guide bias correction is the prediction when the guide scores are set to their median. **b**, CRISPRtile 3D PCA of NLRP3 AlphaFold^1^ structures by alpha carbon where size of the point is the Solvent-Accessible Surface Area where color represents the function score. **c**, cryo-EM structure of the NLRP3 decamer (7PZC)^47^ annotated by the CRISPRtile function score sharing the same colormap as b. **d**, cryo-EM structure of the NLRP3 decamer bound to ADP (7PZC)^47^ annotated by the CRISPRtile function score sharing the same colormap as b. **e**, cryo-EM structure of the NLRP3 decamer bound to the inhibitor MCC950 annotated by the CRISPRtile function score sharing the same colormap as b.

### CRISPRtile enables function-based drug design

Structure-based docking to isolated protein domains is constrained by the assumption that fragments retain their native conformational state. To circumvent the risk of modeling interactions against artificially constrained or misfolded fragments, we used TransformerCPI2.0^48^. Because TransformerCPI2.0^48^ operates independently of 3D structure, it does not force the structural folding of isolated peptides, but uses deep learning to predict compound-protein interactions directly from the sequence^48^. To obtain the functional sequence, CRISPRtile calculates the full width at half maximum (FWHM) of the negative function score relative to the median baseline, obtaining NLRP3 AA62-236. CRISPRtile runs TransformerCPI2.0^48^ on a provided list of compounds or queries FDA approved compounds from ChEMBL^49^, prioritizing candidates predicted to bind to this sequence to induce perturbation or stabilization, with single amino acid deletions across the functional sequence to show stability of the prediction (n=3,229 FDA approved compounds, Fig. 4a).

**Fig. 4.**
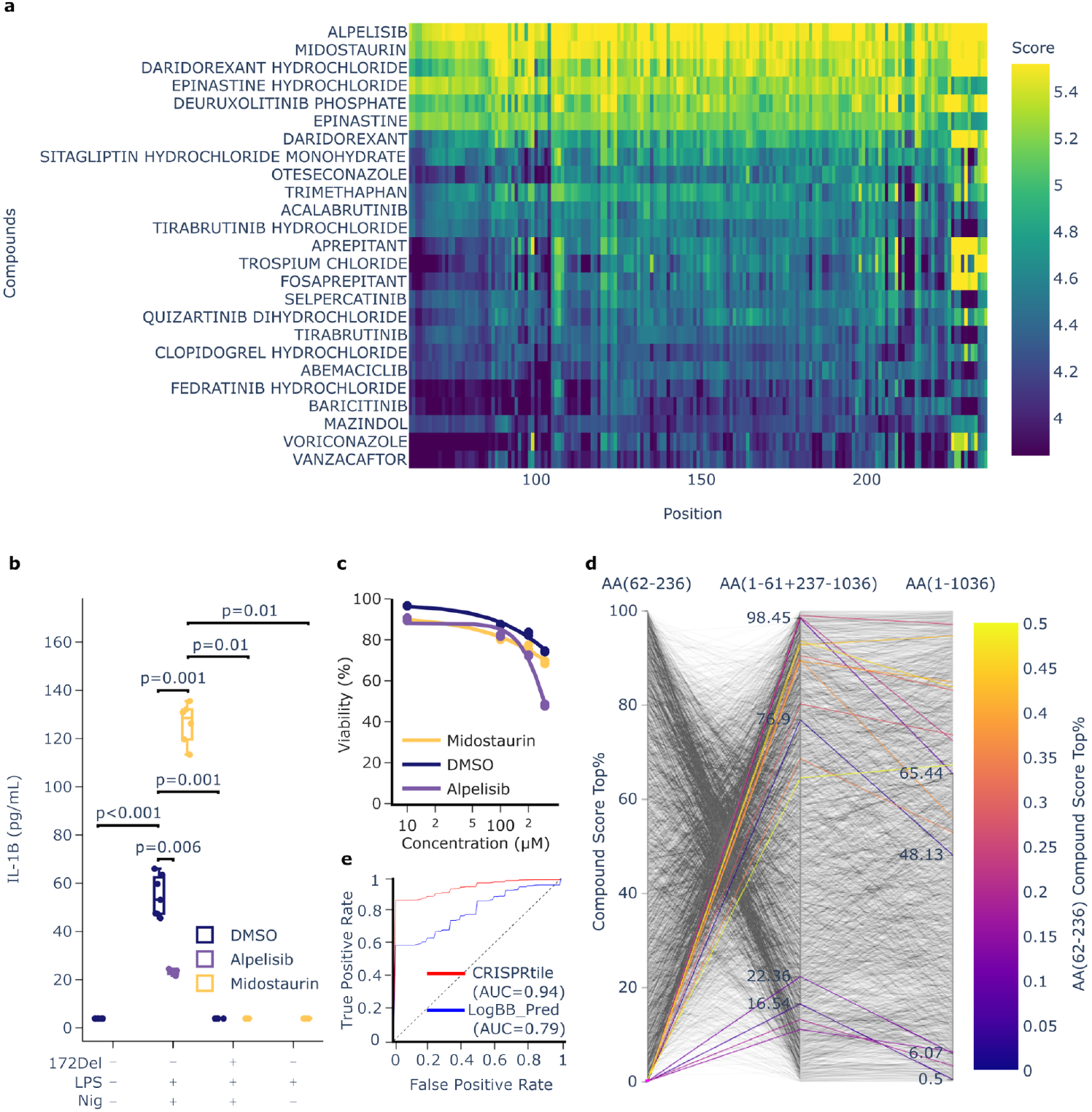
CRISPRtile for function-based drug design. **a**, Logit TransformerCPI2.0^48^ score from amino acid deletions on top 25 scoring compounds across NLRP3 AA62-AA236. **b**, IL-1B ELISA from treatment with compounds. Values under the LOD of 3.9 pg/mL were set to 3.9 pg/mL. DMSO is the control where an equivalent amount is added for compounds resuspended in DMSO. (n = 7, 7, 4, 6, 3, 3, 5 biological replicates for WT+DMSO, WT+LPS+Nig+DMSO, WT+LPS+Nig+Alpelisib, WT+LPS+Nig+Midostaurin, 172del+LPS+Nig+Midostaurin, WT+LPS+Midostaurin, 172del+LPS+Nig+DMSO respectively) P values above the LOD were determined using a two-sided Mann-Whitney U exact test. P values below the LOD were determined using a two-sided fisher exact test. **c**, toxicity dose response curve from flow cytometry live dead staining of THP-1 after treatment with compounds for a day fitted with a 4 parameter logistic regression curve fixed with the minimum value at 0. DMSO is the equivalent amount of DMSO added for the drugs resuspended in DMSO (n = 3 biological replicates) **d**, Comparison of top compound ranking in the functional region identified by CRISPRtile (NLRP3 AA62-236), the functional region removed, and the full protein. **e**, ROC-AUC Curve of CRISPRtile and LogBB_Pred from the mean of bootstrapping 10,000 times on the B3DB^51^ external dataset released after training of the final models, where identical compounds used for training was removed (n = 164 final test set).

We selected the top ranked novel hits, Alpelisib (rank 1) and Midostaurin (Rank 2), and assessed their modulation of IL-1β secretion at a concentration of 10 μM (Fig. 4a-b). This concentration was selected to minimize false positives driven by solubility artifacts (Alpelisib aqueous solubility = 11.96 μM) while maximizing the detection of modulators. Using CRISPRtile’s FACS module for unbiased gating, Midostaurin and Alpelisib had a cell viability of 90.0% and 90.2% respectively, at the selected concentration of 10 μM, ensuring that functional modulations are not a consequence of cytotoxicity (Fig. 4c).

Upon activation in WT, there was increased IL-1β secretion compared to unactivated (p<0.001, Fig. 4b). Alpelisib suppressed the mean IL-1β secretion compared to no Alpelisib by 57.5% (p=0.006, Fig. 4b). Conversely, Midostaurin amplified the mean IL-1β secretion by 131.1% (p=0.001, Fig. 4b). To confirm this amplification was driven by modulation of the NLRP3 pathway, we utilized the genetic ablation of the CRISPRtile predicted functional amino acid (NLRP3 172del) which caused complete loss of detectable IL-1β secretion compared to WT (p=0.001, Fig. 4b). The 172del mutant allows us to differentiate true modulators from artifacts that bypass the NLRP3 pathway. Strikingly, Midostaurin failed to induce detectable secretion in the functional null 172del cells (p=0.01, Fig. 4b), nor did it induce detectable secretion in the absence of the activator Nigericin (p=0.01). This double dependency, requiring both the functional sequence and the upstream trigger, demonstrates that Midostaurin is not an agonist, but a specific amplifier of NLRP3 activation.

We quantified the necessity of the CRISPRtile guided approach by comparing hit rankings against whole protein searches. When the functional sequence was removed, the rankings of our hits dropped precipitously (Fig. 4d). Moreover, in whole protein searches, the compounds lost rank (Midostaurin dropping to the top 0.5% and Alpelisib dropping to the top 48.13%), falling outside standard high throughput screening cutoffs of 0.1-0.3% (Fig. 4d). This demonstrates that without identification of the functional sequence provided by CRISPRtile, these modulators would remain obscured by the global protein surface.

To optimize brain penetration, we integrated a brain penetration prediction model into CRISPRtile. To assess generalization capability beyond the training data, we employed a temporal split validation, where the model was trained exclusively on data available prior to the release of the state-of-the-art benchmark, LogBB_Pred^50^, and evaluated on the subsequently released B3DB^51^ dataset. To preclude data leakage, we removed compounds found in previous datasets used for training. Under these conditions, CRISPRtile achieved an AUC of 0.94, outperforming LogBB_Pred^50^ (AUC=0.79, Fig. 4e). This demonstrates CRISPRtile’s capacity to co-optimize for peripherally restricted or brain penetrant compounds.

## Discussion

CRISPR tiling screens are confounded by guide dependent biases, which can obscure true signals or generate false positives from differences in editing efficiency and indel profile. CRISPRtile overcomes these limitations by integrating AI correction of guide dependent bias with residue specific context. This allows for high resolution sequence-function mapping that would be unattainable with conventional methods. For instance, CRISPRtile successfully unmasked the functional sequence within NLRP3 at positions 62-236 using FWHM of the CRISPRtile function score, a region that existing tools would have discarded. This represents a significant advance over traditional target discovery. Rather than identifying a protein as a target for therapy, CRISPRtile pinpoints the precise sequences required for function, enabling function-based drug design.

Application of CRISPRtile to NLRP3 reveals functional sequences that can be targeted with therapeutic genome editing. Total protein ablation poses risks of toxicity from protein misfolding or double stranded break induced stress^5-7^, and may trigger feedback loops to bring the protein expression back up or remove other functions of the protein required by the cell. In contrast, CRISPRtile identified NLRP3 172 Del, which can be targeted to prevent NLRP3 activation without destabilizing the protein structure leading to the opening of the cage structure^33^ and autoinflammatory disease^15^ (Supplementary Fig. S3b-c). By coupling our high resolution maps with base editing^6,7^, we envision a strategy to perturb these functional sequences. This allows for the therapeutic inhibition of NLRP3 while preserving the protein’s structural integrity and avoiding the deleterious effects of double stranded breaks, offering a safer alternative to broad gene silencing.

We validated the translational utility of CRISPRtile by identifying FDA approved compounds predicted to modulate the functional sequence, allowing us to speed up drug discovery through drug repurposing. We identified Alpelisib as a novel inhibitor suitable for pathologies driven by NLRP3 overactivation, and Midostaurin as a novel amplifier of NLRP3 signaling. The latter offers a potential adjuvant strategy for cancer immunotherapy or infectious disease, enabling localized cytokine release while circumventing the systemic toxicity of cytokine administration. Although CRISPRtile operates independently of 3D structure, docking using DiffDock^56^ (Supplementary Fig. S4) provides a convergent biophysical rationale for these hits. Alpelisib is predicted to occupy the ADP binding site (ΔG≈−7.4 kcal/mol) on the cryo-EM structure (7PZC)^47^, while Midostaurin targets the active conformation (ΔG≈−12.5 kcal/mol) on the cryo-EM structure (8EJ4)^57^. CRISPRtile triangulates functional modulation by converging functional sequence-based prediction, structural docking on functional regions, and live cell functional assessment. By screening on direct modulation of the therapeutic biomarker, CRISPRtile circumvents the reliance on structural surrogates which fail to recapitulate phenotypic outcomes. Furthermore, by integrating pharmacokinetic filters such as brain penetration, CRISPRtile co-optimize for peripherally restricted or brain penetrant compounds.

Ultimately, CRISPRtile establishes a generalizable framework for function-based drug design, transforming vague genetic hits into actionable therapeutic leads. Yet, the translatability of target modulation is constrained in complex polygenic disease or diseases driven by loss of protein function. In scenarios where the protein is truncated, misfolded, or absent, therapeutic efficacy requires direct genome editing to fix the broken component or modulation of compensatory biomarkers to restore homeostatic equilibrium. A promising frontier lies in leveraging genome-wide association studies of disease resilient individuals to pinpoint these targets, subsequently validating their rescue potential via Perturb-seq.

## Acknowledgements

We thank the members of the Center for Translational & Computational Neuroimmunology and the Taub Institute for Research on Alzheimer’s Disease and the Aging Brain for technical assistance and helpful discussions.

We also thank Daniel E. Bauer (Boston Children’s Hospital, Dana-Farber Cancer Institute, Harvard Medical School) and Luca Pinello (Massachusetts General Hospital, Harvard Medical School, Broad Institute) for their valuable input in the development of CRISPRtile.

We acknowledge the CCTI/HICCC Flow Cytometry Core and the resources of the Herbert Irving Comprehensive Cancer Center Flow Cytometry Shared Resource for flow cytometry support. The Flow Cytometry Shared Resource is supported in part by Cancer Center Support Grant P30CA013696.

We also acknowledge the Biomarkers Core Laboratory (BCL) at CUIMC for assistance with ELISA and biochemical assays.

## Materials Availability statement

CRISPR-edited cell lines generated in this study are available from the corresponding author upon reasonable request. The inflammasome modulators used in this study are commercially available (see Methods for details)

## Data Availability

All processed data used is available in Supplementary Tables. The source data is provided with this paper.

## Code Availability

The CRISPRtile platform is available as a Google Colab at (https://colab.research.google.com/drive/14fvBFYtHNU73_JlbyRMZYwv8L84HKisk?usp=sharing). All source code used in this study is available from GitHub at (https://github.com/jasoncngo/CRISPRtile)

## Author Contributions

Conceptualization, Methodology, and Investigation: J.C.N. and F.S.; Data Curation and Formal Analysis: J.C.N., V. A.C.S., X.W., and F.S.; Writing, Review & Editing: J.C.N., X.W., and F.S.; Funding Acquisition and Resources: F.S.; Supervision: F.S.

## Methods

### Synthesis of lentiviral sgRNA libraries

The sgRNA oligos were amplified with PCR using Phusion High-Fidelity DNA Polymerase (NEB, M0530S). The PCR product was then gel purified using QIAquick Gel Extraction Kit (QIAGEN, 28704) and cloned using 10ng of PCR product and 25ng of digested pHKO9 for a 20μl reaction using a Gibson Assembly Master Mix (NEB, E2611S) leaving it at 50C for 60 minutes and 12C after. 1μl of the Gibson Assembly reaction was then added to 25 μl of E.Cloni (Biosearch Technologies, 60107-1) for electroporation using a 1mm Cuvette (Bio-Rad, 165-2089) at 25μF, 200 Ohms, and 1500 volts. The cells were then grown on 24.5 cm bioassay plates coated with LB Agar (Invitrogen, 22700-025) with Ampicillin added. The plasmid library was then extracted from the E. Cloni cells using a NucleoBond Xtra Midi (MACHEREY-NAGEL, 740410.50) and sent for NGS to confirm the distribution of the sgRNAs using a Lorenz curve. To produce the lentivirus, HEK293T cells were cultured with DMEM (Gibco, 11965118) supplemented with 10% fetal bovine serum (Gibco, A5670701) in 15 cm tissue culture dishes. HEK293T cells were then transfected at 70-80% confluence in each plate with 537μl OPTi-MEM (Gibco, 31985070), 135μl Transporter 5 Transfection Reagent (Kyfora Bio, 26008), 9.33μl of 1mg/mL psPAX2 (Addgene #12259), 4.66μl of 1mg/mL VSV-G (Addgene #14888), and 10μg of the plasmid library. The medium was changed 24h after transfection. The lentiviral supernatant was then collected 72h after the transfection and concentrated by ultracentrifugation (26,000 RPM for four hours at 4°C with Beckman Coulter SW 32 Ti rotor). After centrifugation the pellet was resuspended overnight at 4C on a shaker in RPMI+GlutaMAX (Gibco, 61870127) supplemented with 10% fetal bovine serum (FBS) and 2% penicillin-streptomycin (Gibco, 15140122) and stored at -80C.

### Tiled pooled CRISPR-Cas9 screen

LentiCas9-Blast (Addgene #52962) was used to transduce THP-1 cells. THP-1 cells were cultured with RPMI+GlutaMAX supplemented with 10% FBS and 2% penicillin-streptomycin. Transduced cells were then selected using 40μg/ml blasticidin (Invivogen, ant-bl-1). THP-1 cells expressing Cas9 were then transduced at 0.3 MOI with the library lentiviral supernatant while in the expansion phase in 15 cm tissue culture dishes. 24h after the transduction, 40μg/ml blasticidin and 40μg/ml puromycin (Millipore Sigma, P8833) were added to select for cells containing Cas9 and the lentiviral library. 6 and 14 days after transduction, cells were sent for NGS. On the 15th day after transduction, the cells were sorted with FACS.

### ASC speck formation assay

Cells were spun down and resuspended in fresh RPMI+GlutaMAX supplemented with 10% FBS and 2% penicillin-streptomycin containing the cells at a concentration of 500k cells/mL in a 15cm plate with 30mL of media. 12μl LPS (Invitrogen, 00-4976-93) was added to the plate and kept at the 37C incubator for 4 hours. After 4 hours, the cells were then centrifuged and resuspended with 50 ml PBS (Corning, 21-040-CV) that has been preheated to 42C. 75μl of 5mg/ml Nigericin (Millipore Sigma, SML1779-1ML) was added to the resuspended cells in PBS and placed back in the incubator at 42C for 30 min. The cells were then fixed with Cytofix (BD Biosciences, 554714) and left in 4C overnight in the permeabilization medium. The next day, the cells were stained with rabbit polyclonal anti-ASC (AdipoGen, AL177-preservative free) at (1:1000) and kept on ice for 1h. The cells were then washed with the permeabilization medium from Cytofix at 600g for 8 min and incubated on ice for 30 min with the permeabilization medium containing anti-rabbit-IgG-AlexaFluor488 (Invitrogen, A-21206) at (1:750). The cells were then washed and sorted into an Inflammasome activated and unactivated group using the Aria II Cell Sorter with ASC Area vs ASC height. For classification of activated sample ASC speck formation, the CRISPRtile FACS module was used with the false discovery rate set to 0.05.

### Next Generation Sequencing

The cells in the activated and unactivated group post sorting had their genomic DNA extracted using QIAGEN blood and cell culture DNA Midi kit (QIAGEN, 13343) and amplified using the Herculase II PCR (Agilent, 600679). The reaction was performed with 10μl of 5x Herculase II reaction buffer, 0.5μl of 25mM dNTPs, 1μg of genomic DNA, 1.25μl of 10mM forward and reverse primers, 0.5μl of Herculase II fusion DNA polymerase, and water to a total of 50μl for a single reaction and 5 reactions were done for each replicate. The cycling conditions are 98C for 30sec, (98C for 10s, 60C for 20s, and 72C for 20s) done for 23 cycles, and 72C for 2min. The PCR product was then purified using AMPure XP beads (Beckman Coulter, A63881) and a second Herculase II PCR was done using the purified product to add the adapters and indexes. The PCR product was purified again and quantified using KAPA Library Quant Kit (Roche Diagnostics, 07960140001) to send to NGS.

### CRISPRtile Platform

The CRISPRtile FACS module uses XGBoost^52^ with default parameters. For analysis of CRISPR screen, MAGeCK^53^ was used to extract the guide counts. The dropout scores were calculated by log2 fold change in read counts at day 14 over day 6. The function scores were taking the log2 fold change of read counts in the activated population over unactivated population. To ensure robust scoring, scores were calculated for each biological replicate (n = 3) and then averaged to produce a mean score per unique sgRNA. These mean values were used for all downstream statistical comparisons. The predicted double stranded break site of the sgRNAs were then mapped to the protein at the two nearest amino acids. A model was then trained to predict these scores using AutoGluon^54^ with default parameters on the guide dependent scores from CRISPOR^9^, Alphafold^1^ 3D amino acid position, provean score^2^, disorder score, SASA, and amino acid type. MDtraj^37^ was used to obtain SASA and amino acid type. The provean^2^ and disorder scores were obtained from a CRISPRO^36^ reference file. The guide dependent score at all positions was then set to the median guide dependent score to predict the corrected dropout and functional score. The Plotly library in python was used for visualization. The functional sequence was then calculated by CRISPRtile by taking the full width at half maximum of the negative function score relative to the median baseline. TransformerCPI2.0^48^ was run on a list of 3,229 FDA approved compounds queried from ChEMBL^49^ to obtain the compound ranks. The brain penetration prediction model was trained on the B3DB^51^ classification dataset with the exclusion of the external dataset on mordred^55^ molecular descriptors using AutoGluon^54^ for classification and the out-of-fold classification prediction probabilities was used along with the descriptors of the B3DB^51^ LogBB dataset to train the final regression model. DiffDock^56^ was used for structural docking to the cryo-EM structures of activated (8EJ4)^57^ and unactivated (7PZC)^47^ NLRP3. Posebusters^58^ was used on the docked structures for energy minimization. PRODIGY^59^ was used for binding energy predictions.

### Generation of NLRP3 173Del

RNP complex was assembled with 6μl of Cas9 protein (Synthego, #R20SPCAS9-SM), 2.88μl of 50uM guide (IDT), and 1.2μl of AltR E enhancer (IDT, 1075915). The RNP was incubated at room temperature for 20 minutes. THP-1 cells were washed with PBS and resuspended in 18μl of SG cell line solution (Lonza, PBC300675), the RNP complex and 2.4μl of the HDR donor (IDT). The cells were then electroporated with a LONZA 4D Nucleofector unit with electroporation code DV100. The cells were then resuspended in 780μl RPMI+GlutaMAX supplemented with 200μl FBS and 10μl Glutamax (Gibco, 35050-061) and 10μl sodium pyruvate (Gibco, 11360-070) and left in incubator for 72 hours. After 72 hours, cells were plated into 96 well plates at a concentration of 50 cells/plate to isolate single cell clones. After 2 weeks, cells were sent for NGS to confirm editing.

### ELISA

THP-1 cells were seeded at a concentration of 500k cells in 1mL in RPMI+GlutaMAX supplemented with 10% FBS and 2% penicillin-streptomycin. Alpelisib (Santa Cruz Biotechnology, sc391001) and Midostaurin (Santa Cruz Biotechnology, sc-200691) were then resuspended in DMSO (Fisher Bioreagents, BP231-100) and added at a final concentration of 10μM in the media containing the cells at 1μl in 1mL of media. Control cells had DMSO added at a concentration of 1μl in 1mL. 0.4μl of LPS was added concurrently with the drugs and kept at the 37C incubator for 4 hours. 2.5ul of 5mg/ml Nigericin was added to the 1mL of media containing the cells and drugs and kept at the 37C incubator for an hour. The cells were then spun down at 500g for 5min and the supernatant was collected for ELISA using the Human IL-1 beta/IL-1F2 Quantikine ELISA Kit (RND, DLB50).

### Assessment of cell viability with flow cytometry

THP-1 cells were seeded at a concentration of 500k cells in 1mL in RPMI+GlutaMAX supplemented with 10% FBS and 2% penicillin-streptomycin. Cells were treated with Aleplisib and Midostaurin at final concentration of 10μM in the media containing the cells at 1μl in 1mL of media. Control cells had DMSO added at a concentration of 1μl in 1mL. After 24 hours, cells were stained with 7--AAD (BD Biosciences, 559925) at a concentration of 1μg/mL and taken for FACS. Dead cell controls were obtained by treating live cells with 100% DMSO for 15 minutes. For classification of dead cells, the CRISPRtile FACS module was used with the false discovery rate set to 0.05.

### NLRP3 Immunoblot

Protein was harvested from cells using 1x RIPA buffer (Cell Signaling Technology, Cat no. 9806 S) supplemented with protease inhibitor cocktail (RND, 5500) and PMSF (Sigma Aldrich, 93482) and left on ice for an hour. The cell lysate was then centrifuged at 12,000 g for 5 min at 4°C. Half of the supernatant was collected into a separate tube and set on ice. The cell pellet was sonicated and centrifuged at 12,000 g for 5 min at 4°C. The supernatant was then combined with the previously collected supernatant. Protein concentration was quantified using the BCA protein quantification kit(Thermo Scientific, 23225) using a TECAN plate reader. For western blot, 80 µg of protein was loaded into a 4–20% Mini-PROTEAN TGX Precast Protein Gels (Bio-Rad, 4561094). Protein was transferred using Trans-Blot Turbo Mini 0.2 µm PVDF Transfer Packs (BioRad, 1704156). The PVDF membrane was blocked for 40 minutes with a 1:1 ratio of 1x TBS (ThermoFisher, PI28360) and Intercept TBS Blocking Buffer (LICOR, 927-60001). After blocking, 1µg/mL of NLRP3 (Invitrogen, PA579740) and 1:6000 GAPDH (Cell Signaling Technologies, 2118) antibody was added and kept at 4C overnight with gentle shaking. The PVDF membrane was washed three times with 1x TBS-Tween (ThermoFisher Scientific, PI28360) for 8 minutes each. The secondary antibody (LICOR, 92632213) was added and incubated at room temperature for 1 hour with gentle shaking. The PVDF membrane was then washed again three times at 8 minutes each with 1x TBS-Tween. An Odyssey LICOR machine was then used to image the membrane. FIJI ImageJ software was used for quantification.

## References

1. Jumper J, Evans R, Pritzel A, Green T, Figurnov M, Ronneberger O, Tunyasuvunakool K, Bates R, Žídek A, Potapenko A, Bridgland A, Meyer C, Kohl SAA, Ballard AJ, Cowie A, Romera-Paredes B, Nikolov S, Jain R, Adler J, Back T, Petersen S, Reiman D, Clancy E, Zielinski M, Steinegger M, Pacholska M, Berghammer T, Bodenstein S, Silver D, Vinyals O, Senior AW, Kavukcuoglu K, Kohli P, Hassabis D. Highly accurate protein structure prediction with AlphaFold. Nature. 2021 Aug;596(7873):583–589. doi: 10.1038/s41586-021-03819-2. Epub 2021 Jul 15. PMID: 34265844; PMCID: PMC8371605.

2. Choi Y, Sims GE, Murphy S, Miller JR, Chan AP. Predicting the functional effect of amino acid substitutions and indels. PLoS One. 2012;7(10):e46688. doi: 10.1371/journal.pone.0046688. Epub 2012 Oct 8. PMID: 23056405; PMCID: PMC3466303.

3. Sun D, Gao W, Hu H, Zhou S. Why 90% of clinical drug development fails and how to improve it? Acta Pharm Sin B. 2022 Jul;12(7):3049–3062. doi: 10.1016/j.apsb.2022.02.002. Epub 2022 Feb 11. PMID: 35865092; PMCID: PMC9293739.

4. Shi J, Wang E, Milazzo JP, Wang Z, Kinney JB, Vakoc CR. Discovery of cancer drug targets by CRISPR-Cas9 screening of protein domains. Nat Biotechnol. 2015 Jun;33(6):661–7. doi: 10.1038/nbt.3235. Epub 2015 May 11. PMID: 25961408; PMCID: PMC4529991.

5. Anzalone AV, Randolph PB, Davis JR, Sousa AA, Koblan LW, Levy JM, Chen PJ, Wilson C, Newby GA, Raguram A, Liu DR. Search-and-replace genome editing without double-strand breaks or donor DNA. Nature. 2019 Dec;576(7785):149–157. doi: 10.1038/s41586-019-1711-4. Epub 2019 Oct 21. PMID: 31634902; PMCID: PMC6907074.

6. Gaudelli NM, Komor AC, Rees HA, Packer MS, Badran AH, Bryson DI, Liu DR. Programmable base editing of A•T to G•C in genomic DNA without DNA cleavage. Nature. 2017 Nov 23;551(7681):464-471. doi: 10.1038/nature24644. Epub 2017 Oct 25. Erratum in: Nature. 2018 Jul;559(7714):E8. doi: 10.1038/s41586-018-0070-x. PMID: 29160308; PMCID: PMC5726555.

7. Komor AC, Kim YB, Packer MS, Zuris JA, Liu DR. Programmable editing of a target base in genomic DNA without double-stranded DNA cleavage. Nature. 2016 May 19;533(7603):420–4. doi: 10.1038/nature17946. Epub 2016 Apr 20. PMID: 27096365; PMCID: PMC4873371.

8. Concordet JP, Haeussler M. CRISPOR: intuitive guide selection for CRISPR/Cas9 genome editing experiments and screens. Nucleic Acids Res. 2018 Jul 2;46(W1):W242-W245. doi: 10.1093/nar/gky354. PMID: 29762716; PMCID: PMC6030908.

9. Hsu PD, Scott DA, Weinstein JA, Ran FA, Konermann S, Agarwala V, Li Y, Fine EJ, Wu X, Shalem O, Cradick TJ, Marraffini LA, Bao G, Zhang F. DNA targeting specificity of RNA-guided Cas9 nucleases. Nat Biotechnol. 2013 Sep;31(9):827–32. doi: 10.1038/nbt.2647. Epub 2013 Jul 21. PMID: 23873081; PMCID: PMC3969858.

10. Kanduri C, Mamica M, Olstad EW, Zucknick M, Li JJ, Sandve GK. Beware of counter-intuitive levels of false discoveries in datasets with strong intra-correlations. Genome Biol. 2025 Aug 18;26(1):249. doi: 10.1186/s13059-025-03734-z. PMID: 40826107; PMCID: PMC12359981.

11. Cong L, Ran FA, Cox D, Lin S, Barretto R, Habib N, Hsu PD, Wu X, Jiang W, Marraffini LA, Zhang F. Multiplex genome engineering using CRISPR/Cas systems. Science. 2013 Feb 15;339(6121):819–23. doi: 10.1126/science.1231143. Epub 2013 Jan 3. PMID: 23287718; PMCID: PMC3795411.

12. Jinek M, Chylinski K, Fonfara I, Hauer M, Doudna JA, Charpentier E. A programmable dual-RNA-guided DNA endonuclease in adaptive bacterial immunity. Science. 2012 Aug 17;337(6096):816–21. doi: 10.1126/science.1225829. Epub 2012 Jun 28. PMID: 22745249; PMCID: PMC6286148.

13. Canver MC, Smith EC, Sher F, Pinello L, Sanjana NE, Shalem O, Chen DD, Schupp PG, Vinjamur DS, Garcia SP, Luc S, Kurita R, Nakamura Y, Fujiwara Y, Maeda T, Yuan GC, Zhang F, Orkin SH, Bauer DE. BCL11A enhancer dissection by Cas9-mediated in situ saturating mutagenesis. Nature. 2015 Nov 12;527(7577):192–7. doi: 10.1038/nature15521. Epub 2015 Sep 16. PMID: 26375006; PMCID: PMC4644101.

14. Sher F, Hossain M, Seruggia D, Schoonenberg VAC, Yao Q, Cifani P, Dassama LMK, Cole MA, Ren C, Vinjamur DS, Macias-Trevino C, Luk K, McGuckin C, Schupp PG, Canver MC, Kurita R, Nakamura Y, Fujiwara Y, Wolfe SA, Pinello L, Maeda T, Kentsis A, Orkin SH, Bauer DE. Rational targeting of a NuRD subcomplex guided by comprehensive in situ mutagenesis. Nat Genet. 2019 Jul;51(7):1149–1159. doi: 10.1038/s41588-019-0453-4. Epub 2019 Jun 28. PMID: 31253978; PMCID: PMC6650275.

15. Van Gijn ME, Ceccherini I, Shinar Y, Carbo EC, Slofstra M, Arostegui JI, Sarrabay G, Rowczenio D, Omoyımnı E, Balci-Peynircioglu B, Hoffman HM, Milhavet F, Swertz MA, Touitou I. New workflow for classification of genetic variants’ pathogenicity applied to hereditary recurrent fevers by the International Study Group for Systemic Autoinflammatory Diseases (INSAID). J Med Genet. 2018 Aug;55(8):530–537. doi: 10.1136/jmedgenet-2017-105216. Epub 2018 Mar 29. PMID: 29599418.

16. Liston A, Masters SL. Homeostasis-altering molecular processes as mechanisms of inflammasome activation. Nat Rev Immunol. 2017 Mar;17(3):208–214. doi: 10.1038/nri.2016.151. Epub 2017 Feb 6. PMID: 28163301.

17. Lee PH, Yamamoto TN, Gurusamy D, Sukumar M, Yu Z, Hu-Li J, Kawabe T, Gangaplara A, Kishton RJ, Henning AN, Vodnala SK, Germain RN, Paul WE, Restifo NP. Host conditioning with IL-1β improves the antitumor function of adoptively transferred T cells. J Exp Med. 2019 Nov 4;216(11):2619–2634. doi: 10.1084/jem.20181218. Epub 2019 Aug 12. PMID: 31405895; PMCID: PMC6829590.

18. Menachem A, Alteber Z, Cojocaru G, Fridman Kfir T, Blat D, Leiderman O, Galperin M, Sever L, Cohen N, Cohen K, Granit RZ, Vols S, Frenkel M, Soffer L, Meyer K, Menachem K, Galon Tilleman H, Morein D, Borukhov I, Toporik A, Perpinial Shahor M, Tatirovsky E, Mizrachi A, Levy-Barda A, Sadot E, Strenov Y, Eitan R, Jakobson-Setton A, Yanichkin N, Ferre P, Ophir E. Unleashing Natural IL18 Activity Using an Anti-IL18BP Blocker Induces Potent Immune Stimulation and Antitumor Effects. Cancer Immunol Res. 2024 Jun 4;12(6):687–703. doi: 10.1158/2326-6066.CIR-23-0706. PMID: 38592331; PMCID: PMC11148541.

19. Sahoo M, Ceballos-Olvera I, del Barrio L, Re F. Role of the inflammasome, IL-1β, and IL-18 in bacterial infections. ScientificWorldJournal. 2011;11:2037–50. doi: 10.1100/2011/212680. Epub 2011 Nov 1. PMID: 22125454; PMCID: PMC3217589.

20. Kay J, Thadhani E, Samson L, Engelward B. Inflammation-induced DNA damage, mutations and cancer. DNA Repair (Amst). 2019 Nov;83:102673. doi: 10.1016/j.dnarep.2019.102673. Epub 2019 Jul 25. PMID: 31387777; PMCID: PMC6801086.

21. Yang JH, Hayano M, Griffin PT, Amorim JA, Bonkowski MS, Apostolides JK, Salfati EL, Blanchette M, Munding EM, Bhakta M, Chew YC, Guo W, Yang X, Maybury-Lewis S, Tian X, Ross JM, Coppotelli G, Meer MV, Rogers-Hammond R, Vera DL, Lu YR, Pippin JW, Creswell ML, Dou Z, Xu C, Mitchell SJ, Das A, O’Connell BL, Thakur S, Kane AE, Su Q, Mohri Y, Nishimura EK, Schaevitz L, Garg N, Balta AM, Rego MA, Gregory-Ksander M, Jakobs TC, Zhong L, Wakimoto H, El Andari J, Grimm D, Mostoslavsky R, Wagers AJ, Tsubota K, Bonasera SJ, Palmeira CM, Seidman JG, Seidman CE, Wolf NS, Kreiling JA, Sedivy JM, Murphy GF, Green RE, Garcia BA, Berger SL, Oberdoerffer P, Shankland SJ, Gladyshev VN, Ksander BR, Pfenning AR, Rajman LA, Sinclair DA. Loss of epigenetic information as a cause of mammalian aging. Cell. 2023 Jan 19;186(2):305-326.e27. doi: 10.1016/j.cell.2022.12.027. Epub 2023 Jan 12. Erratum in: Cell. 2024 Feb 29;187(5):1312-1313. doi: 10.1016/j.cell.2024.01.049. PMID: 36638792; PMCID: PMC10166133.

22. Ostendorf BN, Bilanovic J, Adaku N, Tafreshian KN, Tavora B, Vaughan RD, Tavazoie SF. Common germline variants of the human APOE gene modulate melanoma progression and survival. Nat Med. 2020 Jul;26(7):1048–1053. doi: 10.1038/s41591-020-0879-3. Epub 2020 May 25. PMID: 32451497; PMCID: PMC8058866.

23. Liu XT, Chen X, Zhao N, Geng F, Zhu MM, Ren QG. Synergism of ApoE4 and systemic infectious burden is mediated by the APOE-NLRP3 axis in Alzheimer’s disease. Psychiatry Clin Neurosci. 2024 Sep;78(9):517–526. doi: 10.1111/pcn.13704. Epub 2024 Jul 16. PMID: 39011734.

24. Fortea J, Pegueroles J, Alcolea D, Belbin O, Dols-Icardo O, Vaqué-Alcázar L, Videla L, Gispert JD, Suárez-Calvet M, Johnson SC, Sperling R, Bejanin A, Lleó A, Montal V. APOE4 homozygozity represents a distinct genetic form of Alzheimer’s disease. Nat Med. 2024 May;30(5):1284–1291. doi: 10.1038/s41591-024-02931-w. Epub 2024 May 6. Erratum in: Nat Med. 2024 Jul;30(7):2093. doi: 10.1038/s41591-024-03127-y. PMID: 38710950.

25. Alijagic A, Hedbrant A, Persson A, Larsson M, Engwall M, Särndahl E. NLRP3 inflammasome as a sensor of micro- and nanoplastics immunotoxicity. Front Immunol. 2023 Apr 18;14:1178434. doi: 10.3389/fimmu.2023.1178434. PMID: 37143682; PMCID: PMC10151538.

26. Nihart AJ, Garcia MA, El Hayek E, Liu R, Olewine M, Kingston JD, Castillo EF, Gullapalli RR, Howard T, Bleske B, Scott J, Gonzalez-Estrella J, Gross JM, Spilde M, Adolphi NL, Gallego DF, Jarrell HS, Dvorscak G, Zuluaga-Ruiz ME, West AB, Campen MJ. Bioaccumulation of microplastics in decedent human brains. Nat Med. 2025 Apr;31(4):1114–1119. doi: 10.1038/s41591-024-03453-1. Epub 2025 Feb 3. Erratum in: Nat Med. 2025 Apr;31(4):1367. doi: 10.1038/s41591-025-03675-x. PMID: 39901044; PMCID: PMC12003191.

27. Coss, R. Could an NLRP3 Inhibitor Be the One Drug to Conquer Common Diseases? C EN Glob. Enterp. 2020, 98, 26–31.

28. Bock C, Datlinger P, Chardon F, Coelho MA, Dong MB, Lawson KA, Lu T, Maroc L, Norman TM, Song B, Stanley G, Chen S, Garnett M, Li W, Moffat J, Qi LS, Shapiro RS, Shendure J, Weissman JS, Zhuang X. High-content CRISPR screening. Nat Rev Methods Primers. 2022;2(1):9. doi: 10.1038/s43586-022-00098-7. Epub 2022 Feb 10. PMID: 37214176; PMCID: PMC10200264.

29. Uijttewaal ECH, Lee J, Sell AC, Botay N, Vainorius G, Novatchkova M, Baar J, Yang J, Potzler T, van der Leij S, Lowden C, Sinner J, Elewaut A, Gavrilovic M, Obenauf A, Schramek D, Elling U. CRISPR-StAR enables high-resolution genetic screening in complex in vivo models. Nat Biotechnol. 2025 Nov;43(11):1848–1860. doi: 10.1038/s41587-024-02512-9. Epub 2024 Dec 16. PMID: 39681701; PMCID: PMC12611787.

30. Petra Berenbrink and Thomas Sauerwald. 2009. The Weighted Coupon Collector’s Problem and Applications. In Proceedings of the 15th Annual International Conference on Computing and Combinatorics (COCOON ‘09). Springer-Verlag, Berlin, Heidelberg, 449–458. 10.1007/978-3-642-02882-3_45

31. Kerur N, Veettil MV, Sharma-Walia N, Bottero V, Sadagopan S, Otageri P, Chandran B. IFI16 acts as a nuclear pathogen sensor to induce the inflammasome in response to Kaposi Sarcoma-associated herpesvirus infection. Cell Host Microbe. 2011 May 19;9(5):363–75. doi: 10.1016/j.chom.2011.04.008. PMID: 21575908; PMCID: PMC3113467.

32. Park YH, Wood G, Kastner DL, Chae JJ. Pyrin inflammasome activation and RhoA signaling in the autoinflammatory diseases FMF and HIDS. Nat Immunol. 2016 Aug;17(8):914–21. doi: 10.1038/ni.3457. Epub 2016 Jun 6. PMID: 27270401; PMCID: PMC4955684.

33. Chae JJ, Park YH, Park C, Hwang IY, Hoffmann P, Kehrl JH, Aksentijevich I, Kastner DL. Connecting two pathways through Ca 2+ signaling: NLRP3 inflammasome activation induced by a hypermorphic PLCG2 mutation. Arthritis Rheumatol. 2015 Feb;67(2):563–7. doi: 10.1002/art.38961. PMID: 25418813; PMCID: PMC4369162.

34. Viganò E, Diamond CE, Spreafico R, Balachander A, Sobota RM, Mortellaro A. Human caspase-4 and caspase-5 regulate the one-step non-canonical inflammasome activation in monocytes. Nat Commun. 2015 Oct 28;6:8761. doi: 10.1038/ncomms9761. PMID: 26508369; PMCID: PMC4640152.

35. Aznauryan E, Yermanos A, Kinzina E, Devaux A, Kapetanovic E, Milanova D, Church GM, Reddy ST. Discovery and validation of human genomic safe harbor sites for gene and cell therapies. Cell Rep Methods. 2022 Jan 14;2(1):100154. doi: 10.1016/j.crmeth.2021.100154. PMID: 35474867; PMCID: PMC9017210.

36. Schoonenberg VAC, Cole MA, Yao Q, Macias-Treviño C, Sher F, Schupp PG, Canver MC, Maeda T, Pinello L, Bauer DE. CRISPRO: identification of functional protein coding sequences based on genome editing dense mutagenesis. Genome Biol. 2018 Oct 19;19(1):169. doi: 10.1186/s13059-018-1563-5. PMID: 30340514; PMCID: PMC6195731.

37. McGibbon RT, Beauchamp KA, Harrigan MP, Klein C, Swails JM, Hernández CX, Schwantes CR, Wang LP, Lane TJ, Pande VS. MDTraj: A Modern Open Library for the Analysis of Molecular Dynamics Trajectories. Biophys J. 2015 Oct 20;109(8):1528–32. doi: 10.1016/j.bpj.2015.08.015. PMID: 26488642; PMCID: PMC4623899.

38. He W, Zhang L, Villarreal OD, Fu R, Bedford E, Dou J, Patel AY, Bedford MT, Shi X, Chen T, Bartholomew B, Xu H. De novo identification of essential protein domains from CRISPR-Cas9 tiling-sgRNA knockout screens. Nat Commun. 2019 Oct 4;10(1):4541. doi: 10.1038/s41467-019-12489-8. Erratum in: Nat Commun. 2020 Feb 25;11(1):1134. doi: 10.1038/s41467-020-14940-7. PMID: 31586052; PMCID: PMC6778102.

39. Apweiler R, Bairoch A, Wu CH, Barker WC, Boeckmann B, Ferro S, Gasteiger E, Huang H, Lopez R, Magrane M, Martin MJ, Natale DA, O’Donovan C, Redaschi N, Yeh LS. UniProt: the Universal Protein knowledgebase. Nucleic Acids Res. 2004 Jan 1;32(Database issue):D115-9. doi: 10.1093/nar/gkh131. PMID: 14681372; PMCID: PMC308865.

40. Oroz J, Barrera-Vilarmau S, Alfonso C, Rivas G, de Alba E. ASC Pyrin Domain Self-associates and Binds NLRP3 Protein Using Equivalent Binding Interfaces. J Biol Chem. 2016 Sep 9;291(37):19487–501. doi: 10.1074/jbc.M116.741082. Epub 2016 Jul 18. PMID: 27432880; PMCID: PMC5016686.

41. Tapia-Abellán A, Angosto-Bazarra D, Alarcón-Vila C, Baños MC, Hafner-Bratkovič I, Oliva B, Pelegrín P. Sensing low intracellular potassium by NLRP3 results in a stable open structure that promotes inflammasome activation. Sci Adv. 2021 Sep 17;7(38):eabf4468. doi: 10.1126/sciadv.abf4468. Epub 2021 Sep 15. PMID: 34524838; PMCID: PMC8443177.

42. Feng S, Wierzbowski MC, Hrovat-Schaale K, Dumortier A, Zhang Y, Zyulina M, Baker PJ, Reygaerts T, Steiner A, De Nardo D, Narayanan DL, Milhavet F, Pinzon-Charry A, Arostegui JI, Khubchandani RP, Geyer M, Boursier G, Masters SL. Mechanisms of NLRP3 activation and inhibition elucidated by functional analysis of disease-associated variants. Nat Immunol. 2025 Mar;26(3):511–523. doi: 10.1038/s41590-025-02088-9. Epub 2025 Feb 10. PMID: 39930093; PMCID: PMC11876074.

43. Hafner-Bratkovič I, Sušjan P, Lainšček D, Tapia-Abellán A, Cerović K, Kadunc L, Angosto-Bazarra D, Pelegrin P, Jerala R. NLRP3 lacking the leucine-rich repeat domain can be fully activated via the canonical inflammasome pathway. Nat Commun. 2018 Dec 5;9(1):5182. doi: 10.1038/s41467-018-07573-4. PMID: 30518920; PMCID: PMC6281599.

44. Landrum MJ, Lee JM, Benson M, Brown GR, Chao C, Chitipiralla S, Gu B, Hart J, Hoffman D, Jang W, Karapetyan K, Katz K, Liu C, Maddipatla Z, Malheiro A, McDaniel K, Ovetsky M, Riley G, Zhou G, Holmes JB, Kattman BL, Maglott DR. ClinVar: improving access to variant interpretations and supporting evidence. Nucleic Acids Res. 2018 Jan 4;46(D1):D1062-D1067. doi: 10.1093/nar/gkx1153. PMID: 29165669; PMCID: PMC5753237.

45. Ohishi A, Masunaga Y, Iijima S, Yamoto K, Kato F, Fukami M, Saitsu H, Ogata T. De novo ZBTB7A variant in a patient with macrocephaly, intellectual disability, and sleep apnea: implications for the phenotypic development in 19p13.3 microdeletions. J Hum Genet. 2020 Jan;65(2):181–186. doi: 10.1038/s10038-019-0690-5. Epub 2019 Oct 23. PMID: 31645653.

46. von der Lippe C, Tveten K, Prescott TE, Holla ØL, Busk ØL, Burke KB, Sansbury FH, Baptista J, Fry AE, Lim D, Jolles S, Evans J, Osio D, Macmillan C, Bruno I, Faletra F, Climent S, Urreitzi R, Hoenicka J, Palau F, Cohen ASA, Engleman K, Zhou D, Amudhavalli SM, Jeanne M, Bonnet-Brilhault F, Lévy J, Drunat S, Derive N, Haug MG, Thorstensen WM. Heterozygous variants in ZBTB7A cause a neurodevelopmental disorder associated with symptomatic overgrowth of pharyngeal lymphoid tissue, macrocephaly, and elevated fetal hemoglobin. Am J Med Genet A. 2022 Jan;188(1):272–282. doi: 10.1002/ajmg.a.62492. Epub 2021 Sep 13. Erratum in: Am J Med Genet A. 2022 Jun;188(6):1930. doi: 10.1002/ajmg.a.62711. PMID: 34515416.

47. Hochheiser IV, Pilsl M, Hagelueken G, Moecking J, Marleaux M, Brinkschulte R, Latz E, Engel C, Geyer M. Structure of the NLRP3 decamer bound to the cytokine release inhibitor CRID3. Nature. 2022 Apr;604(7904):184–189. doi: 10.1038/s41586-022-04467-w. Epub 2022 Feb 3. PMID: 35114687.

48. Chen L, Fan Z, Chang J, Yang R, Hou H, Guo H, Zhang Y, Yang T, Zhou C, Sui Q, Chen Z, Zheng C, Hao X, Zhang K, Cui R, Zhang Z, Ma H, Ding Y, Zhang N, Lu X, Luo X, Jiang H, Zhang S, Zheng M. Sequence-based drug design as a concept in computational drug design. Nat Commun. 2023 Jul 14;14(1):4217. doi: 10.1038/s41467-023-39856-w. PMID: 37452028; PMCID: PMC10349078.

49. Gaulton A, Bellis LJ, Bento AP, Chambers J, Davies M, Hersey A, Light Y, McGlinchey S, Michalovich D, Al-Lazikani B, Overington JP. ChEMBL: a large-scale bioactivity database for drug discovery. Nucleic Acids Res. 2012 Jan;40(Database issue):D1100-7. doi: 10.1093/nar/gkr777. Epub 2011 Sep 23. PMID: 21948594; PMCID: PMC3245175.

50. Shaker B, Lee J, Lee Y, Yu MS, Lee HM, Lee E, Kang HC, Oh KS, Kim HW, Na D. A machine learning-based quantitative model (LogBB_Pred) to predict the blood-brain barrier permeability (logBB value) of drug compounds. Bioinformatics. 2023 Oct 3;39(10):btad577. doi: 10.1093/bioinformatics/btad577. PMID: 37713469; PMCID: PMC10560102.

51. Meng F, Xi Y, Huang J, Ayers PW. A curated diverse molecular database of blood-brain barrier permeability with chemical descriptors. Sci Data. 2021 Oct 29;8(1):289. doi: 10.1038/s41597-021-01069-5. PMID: 34716354; PMCID: PMC8556334.

52. Tianqi Chen and Carlos Guestrin. 2016. XGBoost: A Scalable Tree Boosting System. In Proceedings of the 22nd ACM SIGKDD International Conference on Knowledge Discovery and Data Mining (KDD ‘16). Association for Computing Machinery, New York, NY, USA, 785–794. 10.1145/2939672.2939785

53. Li W, Xu H, Xiao T, Cong L, Love MI, Zhang F, Irizarry RA, Liu JS, Brown M, Liu XS. MAGeCK enables robust identification of essential genes from genome-scale CRISPR/Cas9 knockout screens. Genome Biol. 2014;15(12):554. doi: 10.1186/s13059-014-0554-4. PMID: 25476604; PMCID: PMC4290824.

54. Erickson, Nick, et al. “AutoGluon-Tabular: Robust and Accurate AutoML for Structured Data.” arXiv preprint 2003.06505, 2020.

55. Moriwaki H, Tian YS, Kawashita N, Takagi T. Mordred: a molecular descriptor calculator. J Cheminform. 2018 Feb 6;10(1):4. doi: 10.1186/s13321-018-0258-y. PMID: 29411163; PMCID: PMC5801138.

56. Corso, G., Stärk, H., Jing, B., Barzilay, R. & Jaakkola, T. DiffDock: Diffusion steps, twists, and turns for molecular docking. In Proc. International Conference on Learning Representations (2023).

57. Xiao L, Magupalli VG, Wu H. Cryo-EM structures of the active NLRP3 inflammasome disc. Nature. 2023 Jan;613(7944):595–600. doi: 10.1038/s41586-022-05570-8. Epub 2022 Nov 28. PMID: 36442502; PMCID: PMC10091861.

58. Buttenschoen, M., Morris, G. M. & Deane, C. M. PoseBusters: AI-based docking methods fail to generate physically valid poses or generalise to novel sequences. Chem. Sci. 15, 3130–3151 (2024).

59. Xue LC, Rodrigues JP, Kastritis PL, Bonvin AM, Vangone A. PRODIGY: a web server for predicting the binding affinity of protein-protein complexes. Bioinformatics. 2016 Dec 1;32(23):3676–3678. doi: 10.1093/bioinformatics/btw514. Epub 2016 Aug 8. PMID: 27503228.

