## Supplementary Material for "AI platform for CRISPR functional mapping and function-based drug design"

**Supplementary Figure 1. Raw sorting data of screen.**
**Supplementary Figure 2. NLRP3 172 Del clone and external dataset.**
**Supplementary Figure 3. Additional CRISPRtile outputs.**
**Supplementary Figure 4. Docking of Alpelisib and Midostaurin.**

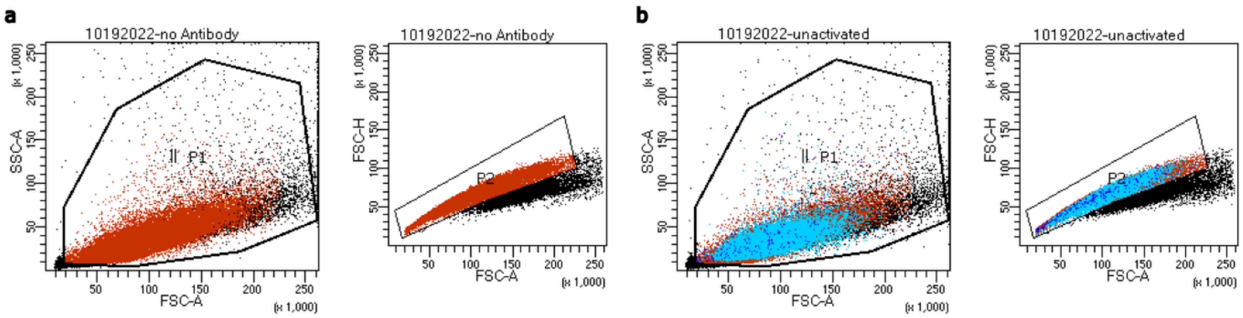


Tube: no Antibody

| Population | #Events | %Parent | %Total |
|---|---|---|---|
| All Events | 1,000,000 | #### | 100.0 |
| P1 | 979,546 | 98.0 | 98.0 |
| P2 | 930,458 | 95.0 | 93.0 |
| Non-Activated | 31 | 0.0 | 0.0 |
| Activated | 1 | 0.0 | 0.0 |

Tube: unactivated

| Population | #Events | %Parent | %Total |
|---|---|---|---|
| All Events | 1,000,000 | #### | 100.0 |
| P1 | 971,514 | 97.2 | 97.2 |
| P2 | 906,749 | 93.3 | 90.7 |
| Non-Activated | 504,189 | 55.6 | 50.4 |
| Activated | 2,485 | 0.3 | 0.2 |

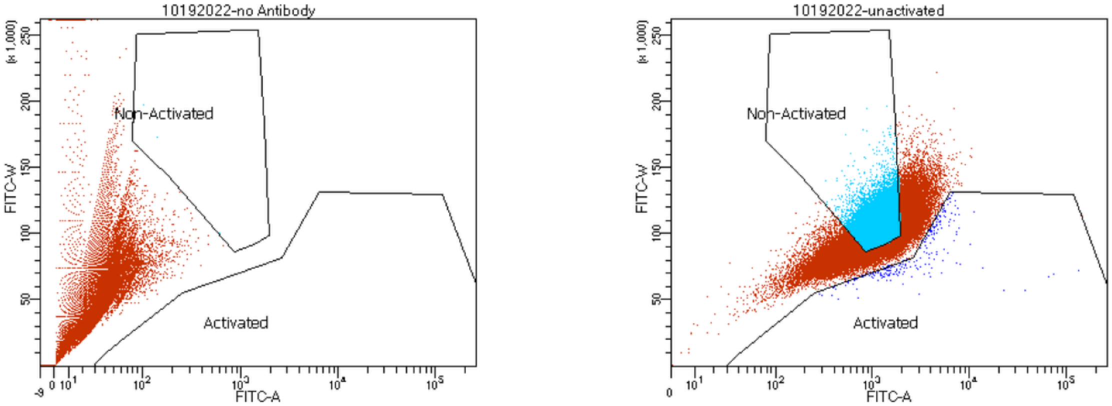


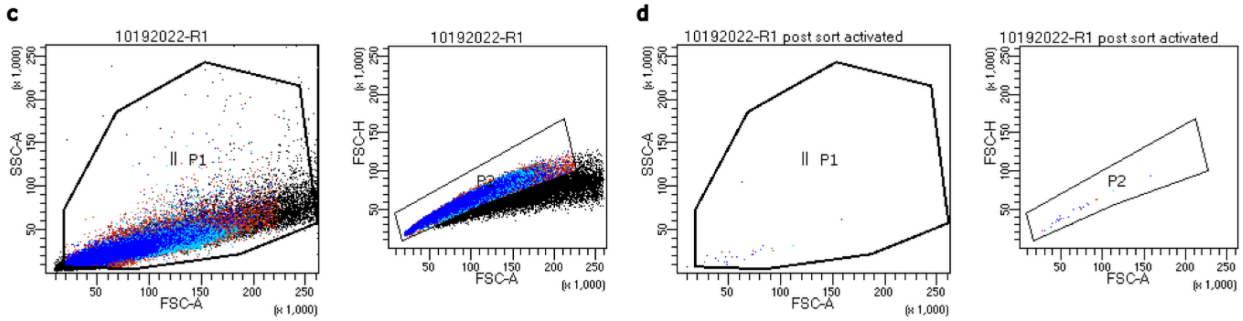


Tube: R1

| Population | #Events | %Parent | %Total |
|---|---|---|---|
| All Events | 1,000,000 | #### | 100.0 |
| P1 | 941,687 | 94.2 | 94.2 |
| P2 | 833,788 | 88.5 | 83.4 |
| Non-Activated | 246,704 | 29.6 | 24.7 |
| Activated | 176,282 | 21.1 | 17.6 |

Tube: R1 post sort activated

| Population | #Events | %Parent | %Total |
|---|---|---|---|
| All Events | 263 | #### | 100.0 |
| P1 | 251 | 95.4 | 95.4 |
| P2 | 250 | 99.6 | 95.1 |
| Non-Activated | 3 | 1.2 | 1.1 |
| Activated | 183 | 73.2 | 69.6 |

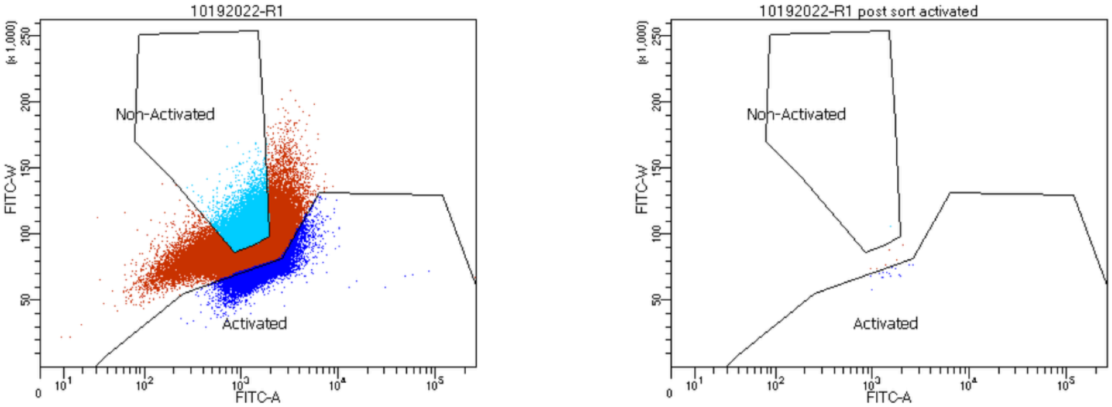

**Supplementary Figure 1. Raw sorting data of screen. a**, FACS data on cells containing the library without primary antibody. **b**, FACS data of cells containing the library **c**, FACS data of cells containing the library after treatment with LPS and Nigericin. **d**, FACS data of cells containing the library sorted in the activated gate and resorted to determine robustness of sort.

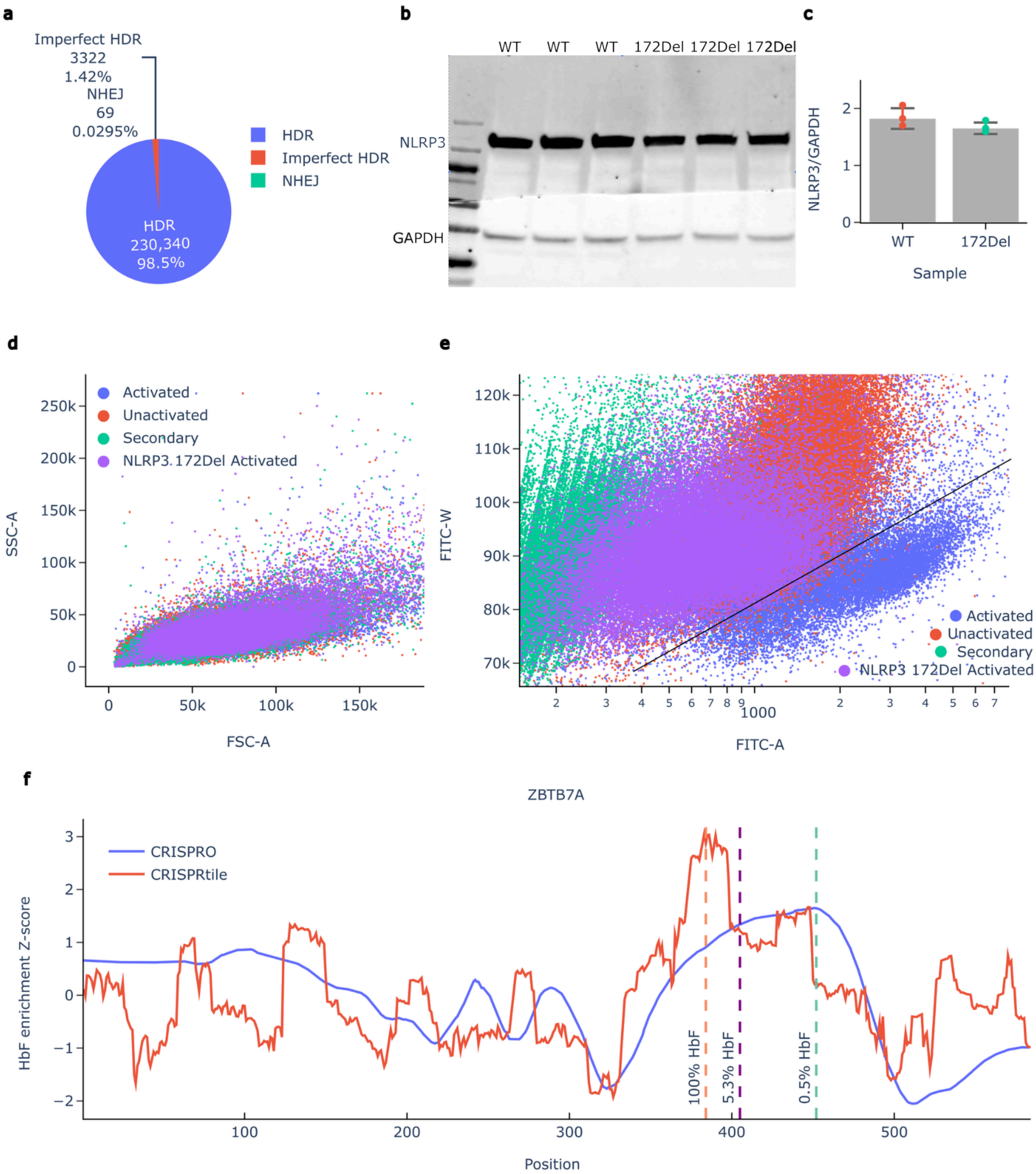


**Supplementary Figure 2. NLRP3 172 Del clone and external dataset. a**, NGS confirmation of HDR editing purity of NLRP3 172 Del clone. **b**, Western blot of WT and NLRP3 172 Del clone for NLRP3 protein expression. **c**, Western blot quantification of WT and NLRP3 172 Del clone for NLRP3 protein expression. **d**, FSC-A and SSC-A of Activated (WT+LPS+Nigericin), Unactivated (WT), Secondary (No primary antibody), and NLRP3 172Del Activated (Clone with LPS+Nigericin) showing that the cells are comparable and results are not driven by cell debris. **e**, FITC-A and FITC-W of Activated (WT+LPS+Nigericin), Unactivated (WT), Secondary (No primary antibody), and NLRP3 172Del Activated (Clone with LPS+Nigericin) showing that NLRP3 172Del Activated does not move into the Activated gate by manual gating indicated by the black line. **f**, CRISPRtile on external dataset tiling ZBTB7A[1] with vertical lines showing clinical variants.

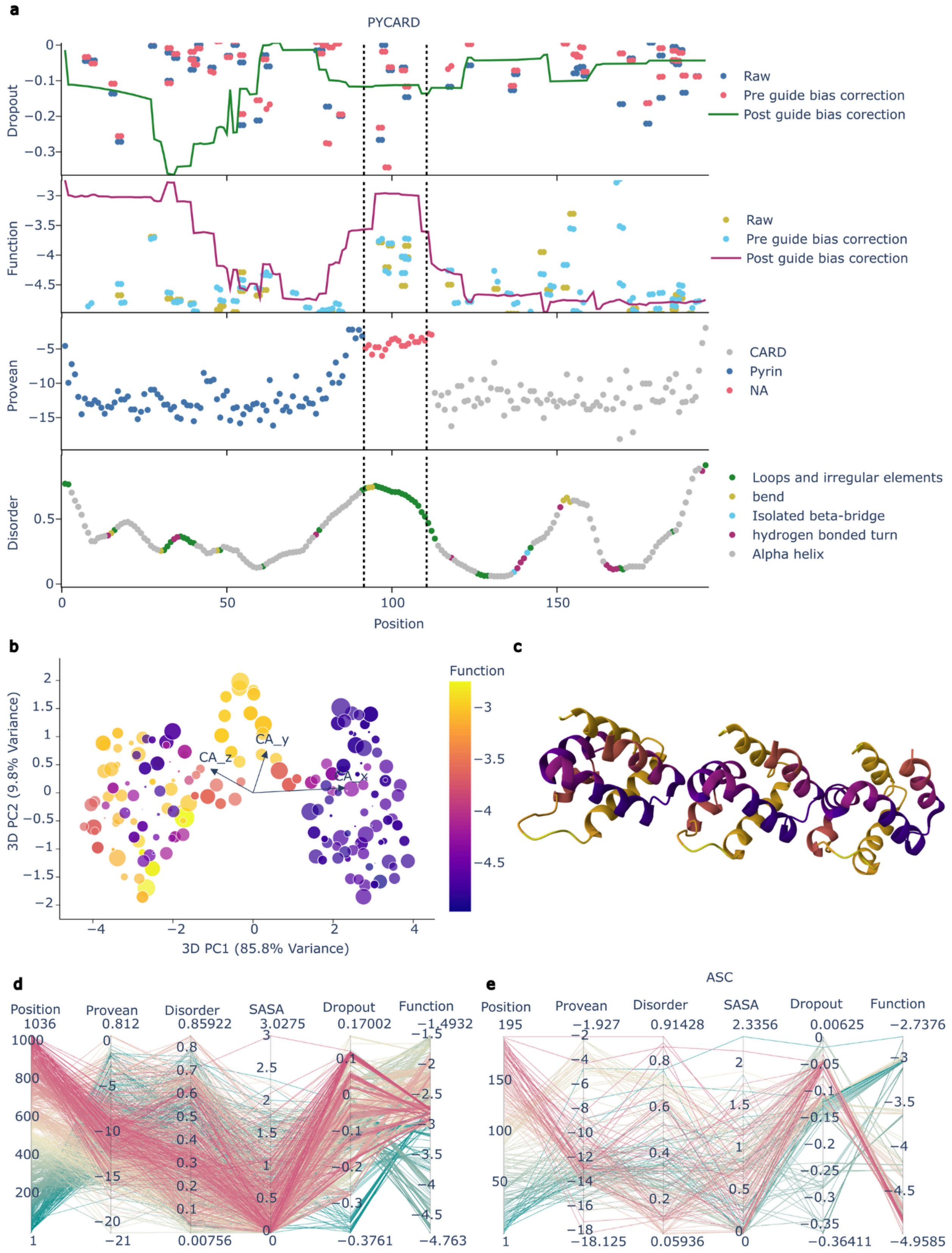


**Supplementary Figure 3. Additional CRISPRtile outputs. a**, CRISPRtile map of

PYCARD by amino acid position where exons are separated by black dotted lines. Raw data points represents the mean log2 fold change across n = 3 biological replicates for unique sgRNAs (ASC, n = 93). The pre guide bias correction is the out-of-fold prediction from 8-fold cross-validation and post guide bias correction is the prediction when the guide scores are set to their median. **b**, CRISPRtile 3D PCA of PYCARD by alpha carbon where size of the point is the Solvent-Accessible Surface Area and color represents the function score. **c**, PYCARD filament from cryo-EM structure (3J63)[2] showing CRISPRtile identified the connection points by functional importance. **d**, Parallel plot of NLRP3 scores. e Parallel plot of PYCARD scores.

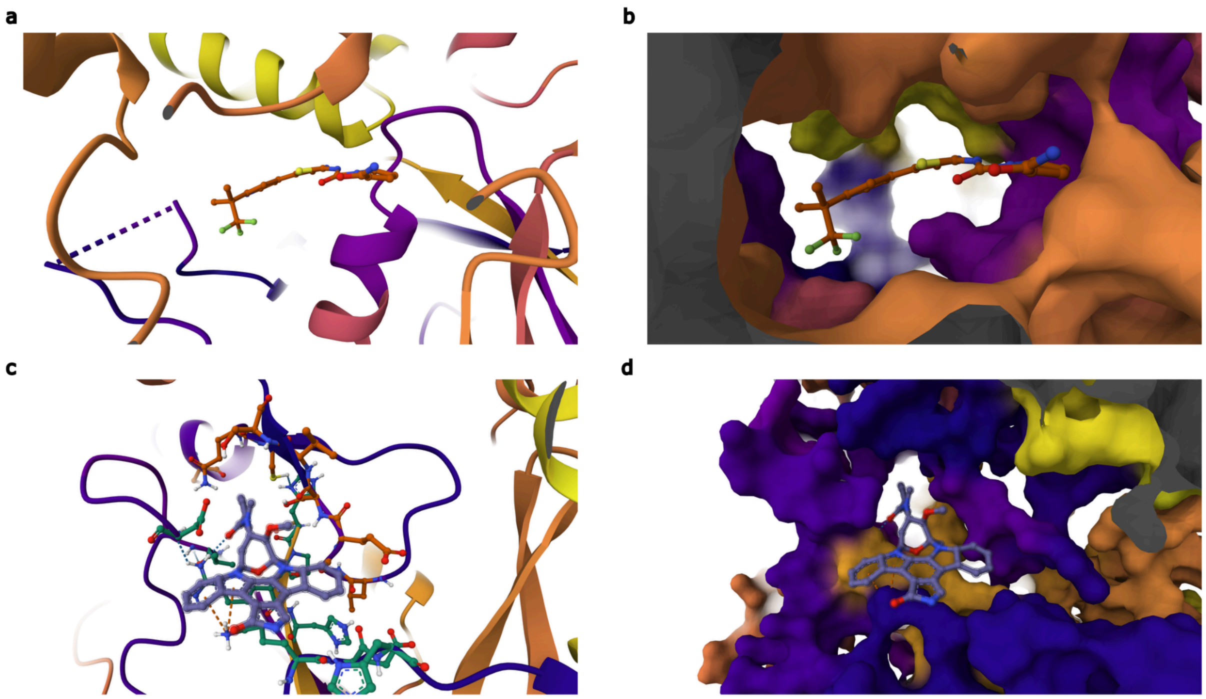


**Supplementary Figure 4. Docking of Alpelisib and Midostaurin. a**, Cartoon of Alpelisib docking to the ADP binding site of the cryo-EM unactivated structure (7PZC)[3]. **b**, Molecular surface of Alpelisib docking to the ADP binding site of the cryo-EM unactivated structure (7PZC)[3]. **c**, Cartoon of Midostaurin docking to the activated NLRP3 structure (8EJ4)[4]. **d**, Molecular surface of Midostaurin docking to the activated NLRP3 structure (8EJ4)[4].

**Supplementary References**


1. Sher F, Hossain M, Seruggia D, Schoonenberg VAC, Yao Q, Cifani P, Dassama LMK, Cole MA, Ren C, Vinjamur DS, Macias-Trevino C, Luk K, McGuckin C, Schupp PG, Canver MC, Kurita R, Nakamura Y, Fujiwara Y, Wolfe SA, Pinello L, Maeda T, Kentsis A, Orkin SH, Bauer DE. Rational targeting of a NuRD subcomplex guided by comprehensive in situ mutagenesis. Nat Genet. 2019 Jul;51(7):1149-1159. doi: 10.1038/s41588-019-0453-4. Epub 2019 Jun 28. PMID: 31253978; PMCID: PMC6650275.
2. Lu A, Magupalli VG, Ruan J, Yin Q, Atianand MK, Vos MR, Schröder GF, Fitzgerald KA, Wu H, Egelman EH. Unified polymerization mechanism for the assembly of ASC-dependent inflammasomes. Cell. 2014 Mar 13;156(6):1193-1206. doi: 10.1016/j.cell.2014.02.008. PMID: 24630722; PMCID: PMC4000066.
3. Hochheiser IV, Pilsl M, Hagelueken G, Moecking J, Marleaux M, Brinkschulte R, Latz E, Engel C, Geyer M. Structure of the NLRP3 decamer bound to the cytokine release inhibitor CRID3. Nature. 2022 Apr;604(7904):184-189. doi: 10.1038/s41586-022-04467-w. Epub 2022 Feb 3. PMID: 35114687.
4. Xiao L, Magupalli VG, Wu H. Cryo-EM structures of the active NLRP3 inflammasome disc. Nature. 2023 Jan;613(7944):595-600. doi: 10.1038/s41586-022-05570-8. Epub 2022 Nov 28. PMID: 36442502; PMCID: PMC10091861.
